# CPT1B-Mediated Fatty Acid Oxidation Induces Pigmentation in Solar Lentigo

**DOI:** 10.1101/2025.06.15.659823

**Authors:** Yueun Choi, Uijeong Nam, Jihan Kim, Soo-Young Yoon, Nahyun Lee, Hanseul Cho, Seoyeon Kim, Sanghyeon Yu, Junghyun Kim, Hae-Na Moon, Bark-Lynn Lew, Yoonsung Lee, Man S Kim, Soon-Hyo Kwon

## Abstract

**Introduction:** Cellular senescence is associated with altered lipid metabolism, including increased cellular lipid uptake, upregulated lipid biosynthesis, and deregulated lipid breakdown. Previous studies have reported that carnitine palmitoyltransferase (CPT), the rate-limiting enzyme in fatty acid oxidation that catalyzes the conversion of acyl-CoA to acylcarnitine, is involved in various senescence-related diseases. Although solar lentigo (SL) is an age-related pigmentary disorder characterized by the accumulation of senescent cells, its role in metabolic dysregulation has rarely been investigated.

**Methods and Results:** Integrated transcriptomic profiling of SL skin samples, combining mRNA sequencing, differential gene expression, pathway enrichment analyses, metabolic flux simulations, and protein-protein interaction analysis, was conducted to demonstrate the molecular alterations in SL compared to perilesional normal skin. We found transcriptomic alterations in mitochondrial energy metabolism-associated genes. Metabolic flux simulations revealed that carnitine-associated reactions involved in fatty acid oxidation were upregulated. Using a multi-omics approach, *CPT1B* was selected as a potential marker for SL, which was confirmed via its overexpression in immunohistochemical studies. Using a zebrafish model, *CPT1B* was implicated in melanogenesis.

**Discussion:** *CPT1B*-mediated metabolic alteration is a key driver of SL pathogenesis. Targeting *CPT1B* and the associated lipid metabolism pathways is a novel therapeutic approach for managing SL and age-related pigmentation disorders.

**Research Highlights:** Solar lentigo (SL) exhibits altered mitochondrial metabolism and fatty acid oxidation. *CPT1B* activation regulates pigmentation, suggesting that targeting *CPT1B* may offer a novel therapy for SL and other age-related pigmentary disorders.

**Graphical abstract:** 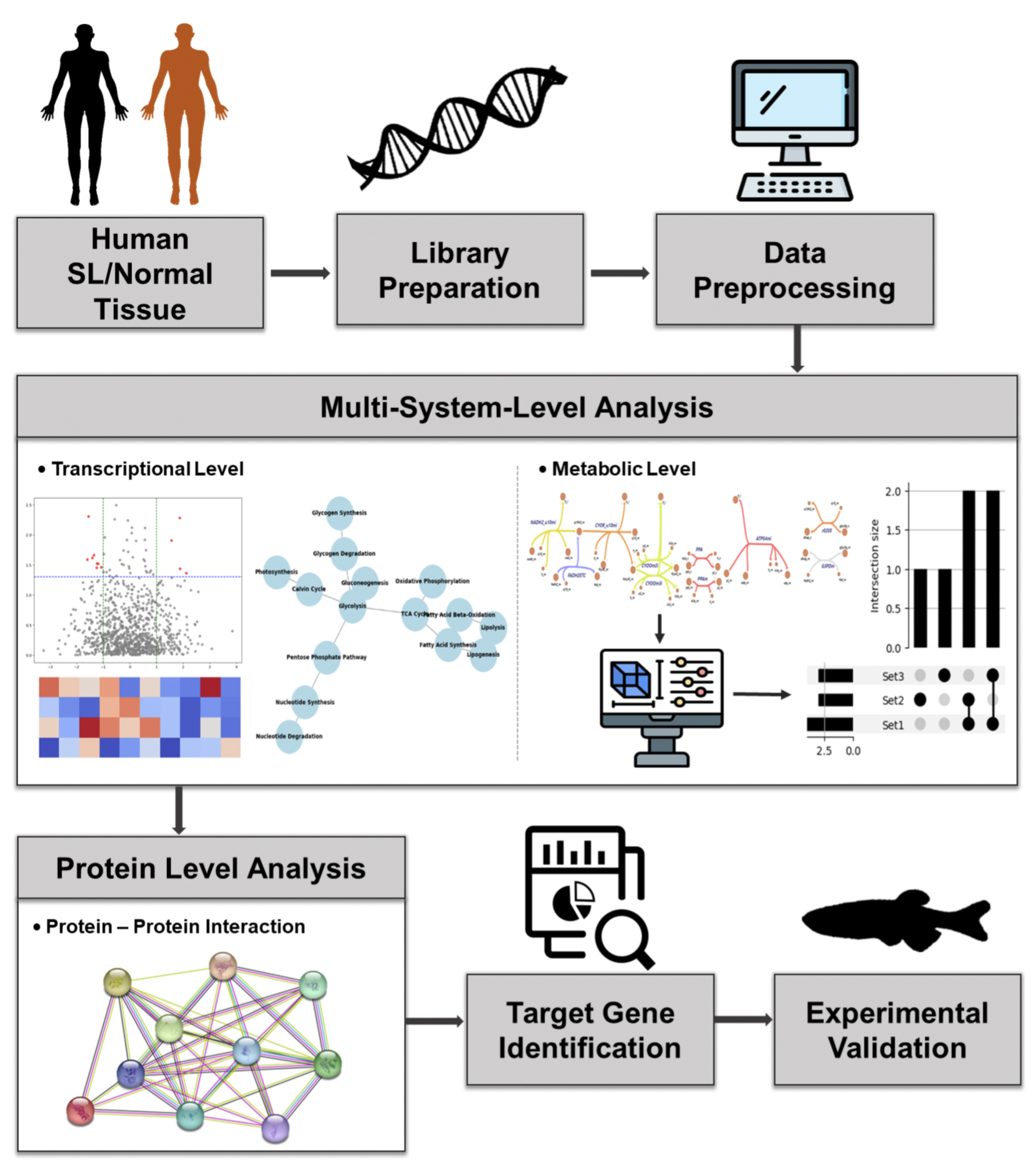

## Introduction

Aging is a functional decline that occurs during adulthood. The hallmarks of aging include cellular senescence, mitochondrial dysfunction, deregulated nutrient sensing, genomic instability, telomere attrition, epigenetic alterations, loss of proteostasis, disabled macroautophagy, stem cell exhaustion, altered intercellular communication, chronic inflammation, and dysbiosis, which occur in close connection with each other (Lopez-Otin et al., 2023). For example, cellular senescence, which is induced in part by telomere shortening with ageing, plays a pathologic role in age-related diseases via the tissue-specific accumulation of senescent cells and secretion of senescence-associated secretory phenotypes (SASPs)(Blasco, 2005). Cellular senescence also affects the metabolic process of cells, with a marked increase in mitochondrial activity observed in senescent cells (Correia-Melo et al., 2016). While young fibroblasts have glycolytic phenotype and utilize few free fatty acids and amino acids, aged fibroblasts have increased secretion of lipids and high expression of fatty acid synthase (Alicea et al., 2020; Kruglikov et al., 2022). Moreover, metabolic alteration can affect the cells by determining senescence entry. Inhibition of fatty acid synthase prevents the induction of senescence in human fibroblasts, resulting in reduced *p*53-induced senescence and decreased SASP expression (Borghesan et al., 2019; Fafian-Labora et al., 2019). Defects in cellular metabolism or the distribution of mitochondria into daughter cells can modulate stem cell populations towards exhaustion (Ito & Ito, 2016). Collectively, cellular senescence is closely associated with mitochondrial dysfunction and metabolic alteration.

Solar lentigo (senile lentigo, SL) is a common acquired pigmentary disorder considered a hallmark of skin aging (Jeong et al., 2024). Senescent keratinocytes with enlarged morphology are observed in SL, and these could contribute to the thickened epidermis observed (Lin et al., 2010; Shin et al., 2015). They also exhibit decreased proliferation, impaired differentiation, and hyperplasia, which are implicated in cellular senescence (Lin et al., 2010; Shin et al., 2015). *p*16^INK4A^-positive senescent fibroblasts are more prevalent in the upper dermis than in the normal perilesional skin (Yoon et al., 2018). Senescent fibroblasts produce higher levels of pro-melanogenic stimuli, including hepatocyte growth factor, keratinocyte growth factor, and stem cell factor than young fibroblasts, resulting in the upregulated melanogenesis (Hasegawa et al., 2015; Kovacs et al., 2010; Lin et al., 2010; Shin et al., 2012). Recently, several SASPs from senescent fibroblasts, such as stromal cell-derived factor-1 and growth/differentiation factor-15, have induced pigmentation in SL via epigenetic change or β-catenin signaling (Kim et al., 2020; Yoon et al., 2018). Although increased expression of fatty acid metabolism-related genes has been reported, a comprehensive analysis covering transcriptional and metabolic alterations involved in SL development has never been investigated (Aoki et al., 2007). This study aimed to elucidate the transcriptional-metabolic alterations implicated in SL development by identifying the corresponding regulatory mechanisms of differentially activated metabolic reactions and their potential enzymatic genes at the multisystem level. To investigate the specific underlying gene regulation, we obtained transcription profiles from SL skin samples using mRNA sequencing, and utilized several comparative analytical approaches and an in silico model. Differential analyses comparing SL and perilesional normal skin at the transcriptional level were conducted using three distinct gene lists: differentially expressed genes (DEGs), a customized mitochondria-associated gene group, and MitoCarta3.0. Pathway enrichment analyses at the transcriptional level, such as gene set enrichment analysis (GSEA) and fast GSEA (FGSEA), were performed for comparison. Additionally, metabolic flux simulations were performed to identify potential metabolic reactions that were significantly altered during SL development. Given the substantial alterations observed at the transcriptional and metabolic levels, a potential target gene list was curated. Target genes were validated using immunohistochemistry and a zebrafish model.

## Materials and Methods

### Sample collection

Paired skin samples (5 mm diameter) were obtained from the SL and adjacent normal skin of seven patients and used for mRNA sequencing. This study was conducted in accordance with the Declaration of Helsinki and the International Conference on Harmonization and Good Clinical Practice Guidelines, and was reviewed and approved by the Institutional Review Board of Kyung Hee University Hospital at Gangdong (KHNMC 2022-04-014). Written informed consent was obtained from all participants prior to their enrolment in the study.

### Library construction

The total RNA concentration was measured using the Quant-IT RiboGreen assay (Invitrogen). The RNA integrity was assessed by determining the DV200 value, which indicates the percentage of RNA fragments longer than 200 bp, using the TapeStation RNA Screentape System (Agilent). For sequencing library construction, 100 ng of total RNA was processed using an Agilent SureSelect RNA Direct kit according to the manufacturer’s instructions. Initially, total RNA was fragmented into smaller pieces using divalent cations at elevated temperatures. The fragmented RNA was reverse-transcribed into first-strand cDNA using random primers, followed by second-strand cDNA synthesis. Post-cDNA synthesis, the fragments were subjected to end repair, addition of a single adenine (A) base, and adapter ligation. The products were subsequently purified and enriched using PCR to form a cDNA library. An Agilent SureSelect XT Mouse All Exon Kit was used to capture human exonic regions following the Agilent SureSelect Target Enrichment protocol. Approximately 250 ng of the cDNA library was mixed with hybridization buffers, blocking mixes, RNase block, and 5 µL of the SureSelect All Exon capture library. Capture baits were hybridized at 65°C with a heated thermal cycler lid set to 105°C for 24 h, using a PCR machine. After hybridization, the captured library was washed and subjected to a second round of PCR amplification. The final purified product was quantified using the KAPA Library Quantification Kit for Illumina sequencing platforms, following the qPCR Quantification Protocol Guide (KAPA BIOSYSTEMS, #KK4854), and qualified using TapeStation D1000 ScreenTape (Agilent Technologies, #5067-5582). The indexed libraries were sequenced on an Illumina NovaSeq platform (Illumina, Inc., San Diego, CA, USA) using paired-end sequencing (2 × 100 bp).

### Data processing

Paired-end sequencing reads were generated using the Illumina NovaSeq sequencing platform. Before analysis, Trimmomatic v0.38 was used to remove adapter sequences and trim low-quality bases. The cleaned reads were aligned to the human reference genome (GRCh38) using HISAT v2.1.0 (Kim et al., 2015) and Bowtie2 implementations. Reference genome sequences and gene annotation data were obtained from the NCBI genome assembly and RefSeq databases, respectively. The aligned data in the SAM file format were sorted and indexed using SAMtools v1.9. Post-alignment, transcripts were assembled and quantified using StringTie v2.1.3b (Pertea et al., 2016; Pertea et al., 2015). Gene-level and transcript-level quantifications were acquired as raw read counts and transcripts per million (TPM).

### Differential and enrichment analysis

Differential expression analysis between the SL and control samples was performed using the DESeq2 R package (version 1.38.3) based on TPM levels. Genes were collected by searching the keyword ‘skin’ in the MSigDB and curating ontology gene sets for Homo sapiens across all contributors. A volcano plot was generated to visualize DEGs using the EnhancedVolcano R package (version 1.16.0). For gene ontology analysis, the enrichGO function from the ClusterProfiler R package (version 4.6.2) was used to identify enriched pathways in the SL and control tissues separately. Pathway associations were examined using GSEA (version 4.3.2), and network visualization was performed using the EnrichmentMap application in Cytoscape. Further analysis was performed using FGSEA (version 1.24.0) to compute the normalized enrichment scores for gene modules from MitoCarta 3.0 gene list. Additionally, heatmap analysis of the gene expression profiles was performed using a customized metabolism-associated gene list (Guarnieri et al., 2023).

### Metabolic flux simulation

We used a custom-made context-specific constraint-based metabolic modelling approach,(Camera et al., 2024) which was adapted and refined from previous studies (da Silveira et al., 2020; Guarnieri et al., 2023). This approach was constructed based on the RECON3D metabolic model(Brunk et al., 2018) using CORDA (Schultz & Qutub, 2016) and implemented through Cobrapy (Ebrahim et al., 2013), incorporating gene-reaction rules where enzyme expression levels were linked to their corresponding metabolic reactions. Nine essential metabolic pathways, including ‘oxidative phosphorylation (OXPHOS)’, ‘citric acid cycle’, ‘glycolysis/gluconeogenesis’, ‘CoA synthesis’, ‘CoA catabolism’, ‘NAD metabolism’, ‘fatty acid synthesis’, ‘fatty acid oxidation’, and ‘biomass and maintenance functions’, were manually activated to ensure the stability of the simulation model. All Recon3D reactions were iteratively optimized using the Gurobi solver to compute the flux levels of the corresponding reactions.

Although gene expression levels were incorporated into the model for each sample, all other parameters were kept identical across all simulations. The simulation outcomes were reported as flux levels of all available reactions through custom-made flux balance analysis, where the corresponding levels were analyzed as grouped variables to compare the ‘SL’ and ‘control’ groups. Owing to the difficulty in assuming variance or normality between ‘SL’ and ‘controls’, a non-parametric van der Waerden test was performed to appropriately compare the flux levels, using the R matrixTests package (v. 0.1.9). Metabolic flux alteration was visualized through upset plot, heatmap, and Escher (King et al., 2015).

### Immunohistochemical analysis

Paired skin samples (diameter: 3 mm) of the SL and adjacent normal skin from two female patients (aged 81 years) were obtained for immunohistochemical analysis. Tissue samples were fixed with 10% formalin, embedded in paraffin, and sectioned at 3 μm thickness. Sections were deparaffinized and subjected to antigen retrieval by boiling them in citrate buffer (pH 6.0) at 95°C for 10 min. To reduce nonspecific background staining, the sections were incubated with Ultra V Block (Thermo Scientific, TA-060-UB) for 5 min at room temperature. The sections were incubated overnight with primary anti-CPT1b, 1:200 (ab134988, Abcam). The slides were washed with PBS and incubated with a biotinylated secondary antibody (Polink2 plus HRP Rabbit Dab kit; Origene D41-125, WA, USA) for 30 min at room temperature. After washing with PBS, the peroxidase activity was developed with 3, 3’-diaminobenzidine (DAB, ab64238). Sections were counterstained with hematoxylin (ab220365) and imaged.

### Zebrafish strains

The zebrafish used in our experiments were the wild-type TAB5 strain, raised and maintained at Kyung Hee University Hospital in Gangdong, in accordance with the Institutional Animal Care and Use Committees (IACUC: KHNMC AP 2023). The embryos were raised in E3 solution at 28℃.

### Guide RNA synthesis and Cas9 ribonucleoprotein (RNP) complex preparation

The template sequence for generating *cpt1b*-targeting guide RNA was prepared following the protocol described by Wu et al (Wu et al., 2018). The guide RNA was synthesized *in vitro* using the HiScribe® T7 High Yield RNA Synthesis Kit (NEB) and subsequently purified with the RNA Clean & Concentrator Kits (Zymo Research). Finally, the synthesized guide RNA was combined with Cas9 protein (IDT) and incubated at 37℃ for 5 min to form the RNP complex.

### Microinjections of morpholino (MO), mRNA, and Guide RNA/Cas9

Microinjections were performed using a FemtoJet 4i microinjector (Eppendorf) with borosilicate glass needles prepared using P-1000 next-generation micropipette pullers (Sutter Instruments). Approximately 1 nL of MO, mRNA, or guide RNA/Cas9 RNP was injected into the yolk at the single-cell stage. The concentration of the materials injected was as follows: 5 ng/nL of splicing-blocking cpt1b MO with the sequence 5’-ATCTGCTGTTTAATTTACCTCACTGG-3’; 200 pg/nL of mRNA; and for the guide RNA/Cas9 RNP, 50 pg of each of four guide RNAs targeting *cpt1b* (totaling 200 pg), combined with 800 pg of Cas9 protein.

### Genotyping

Whole embryos at 3 days post fertilization (dpf) were lysed by adding 25 μL of extraction solution and 7 μL tissue preparation solution (Sigma-Aldrich), followed by incubation at 55℃ for 10 min and 95℃ for 20 min. Subsequently, 25 μL neutralization solution (Sigma-Aldrich) was added to neutralize the mixture. The extracted genomic DNA was amplified by PCR, targeting specific regions using the designed primers (primer set: *cpt1b* t1_forward: 5’-TGTTTACCCTGCCAGTCCTT-3’, *cpt1b* t1_reverse: 5’-GCCCAATCAGTACCGAGAAC-3’, *cpt1b* t2_forward: 5’-TTGAGTGCCATCCTGTTTGC-3’, *cpt1b* t2_reverse: 5’-CGACCAGACAGCAGCTTTAC-3’, *cpt1b* t3, 4_forward: 5’-TAACACAGCCAAACGGTGAC-3’, *cpt1b* t3, 4_reverse: 5’-AAGAGCACATGGGGACAATC-3’). Sequencing of PCR amplicons revealed mutations close to the target sites in *cpt1b* crispants.

### RT-PCR

To validate the efficiency of the cpt1b MO, RT-PCR analysis was performed. RNA was extracted using TRIzol™ Reagent (Invitrogen), treated with DNase1 (NEB) to remove any contaminating DNA, and then reverse-transcribed into cDNA using the SuperScript™ IV First-Strand Synthesis System (Invitrogen). To specifically amplify the region targeted by MO, the following primer set was used: cpt1b MO_forward: 5’-CAGTTCACCGTCACTCCTGA-3’ and *cpt1b* MO_reverse: 5’-CGCAGCAGGAATATGACAGA-3’. The resulting PCR amplicons were separated on a 1% agarose gel and visualized using the iBright™ CL1500 Imaging System (Invitrogen).

### Microscopy and image analysis

All images of live embryos used in this study were captured using a Leica M205 FCA microscope (Leica Microsystems).

## Results

### Clinical and histological characteristics of SL sample and its transcriptomic analysis revealing differentially expressed genes

Paired skin samples were obtained from the SL and adjacent normal skin of seven patients (six females and one male). The mean age was 82.0 ± 10.2 years (range, 67–96 years). The location on the chin (71.4%), cheek (14.3%), and temple (14.3%). Histologically, basal hyperpigmentation and solar elastosis were present in all the SL samples, whereas dermal melanophages (71.4%) and rete ridge elongation (28.5%) were also observed. Inflammatory features, including infiltration of inflammatory cells (85.7%) and interface changes (28.5%), were of mild grade.

Using mRNA sequencing data, multiple bioinformatics packages were used for data preprocessing to facilitate downstream analyses. To explore the basic information for RNA-Seq, we applied typical differential analyses to the gene expression profiles. **Supplementary Figure S1A** presents a volcano plot, with red dots denoting significantly activated or inhibited genes in the SL, compared with those in the control. The number of DEGs was sufficient for subsequent downstream analyses. **Supplementary Figure S1B** presents dot plots highlighting the activated or inhibited pathways in SL. Pathways involved in epidermal development and differentiation were enriched in the SL.

### SL exhibits alterations in mitochondria-associated metabolic pathways

To elucidate the core functional mechanisms of SL at the molecular and cellular levels, we focused on metabolic processes, as we discovered metabolic dysregulation using analytical approaches such as GSEA. As shown in **Figure 1A**, while revealing several dysregulated clusters in SL compared to the control, GSEA revealed the largest cluster, negative regulation of phosphate metabolic processes. This finding is consistent with that of a previous report, suggesting that pathways involving protein phosphorylation contribute to the hyperpigmentation observed in the SL (Kadono et al., 2001).

**Figure 1.**
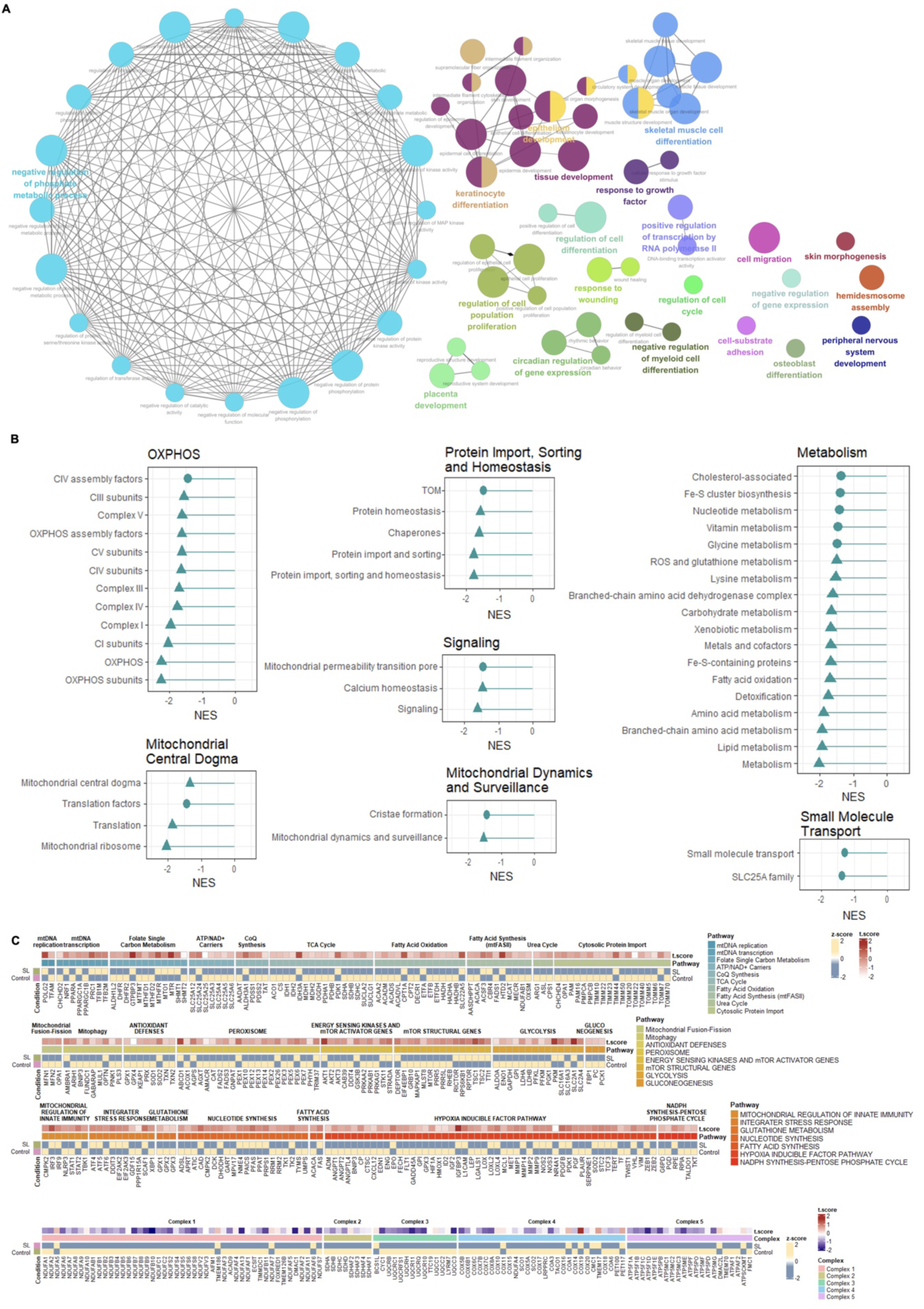
Functional enrichment analysis results based on gene set enrichment analysis (GSEA) and MitoCarta3.0 gene set (A) Network visualization depicting interconnections between selected gene ontology pathways enriched in the differential expression analysis comparing solar lentigo and normal tissue. (B) Lollipop plots displaying normalized enrichment scores of MitoCarta3.0 gene set computed through the fast GSEA R package. Circle dots indicated pathways with *P* < 0.05, while triangle dots represented pathways with *P ≥* 0.05 and < 0.1. (C) Heatmap visualizing t-scores and expression profiles of genes within curated mitochondrial energy metabolism-associated gene sets. SL, solar lentigo.

Recognizing that metabolic dysregulation could be a significant feature of SL, we further analyzed these changes by estimating alterations in energy metabolism-associated pathways that are essential for metabolic processes. In this analysis, we used FGSEA, an appropriate tool for evaluating the dynamics of specific gene groups (i.e., pathways or modules). The MitoCarta3.0 gene list, which encompasses genes associated with energy metabolism, was utilized in the analysis. The results showed systematic downregulation of mitochondria-associated metabolic pathways, including ‘OXPHOS’, ‘mitochondrial central dogma’, ‘mitochondrial dynamics and surveillance’, ‘protein import, sorting and homeostasis’, and ‘signaling’ (**Figure 1B**). This observation is congruent with previous studies that demonstrated a significantly higher incidence of the common 4977 bp deletion in mitochondrial DNA in 50% of SL than in adjacent normal skin (Hafner et al., 2011). This common deletion is present in ageing skin, particularly in chronologically aged and photoaged skin.

### SL exhibits upregulated expression of the specific genes involved in mitochondrial energy metabolism-associated pathways

Considering that the molecular activities associated with pivotal mechanisms in SL are likely to exhibit higher expression levels at the transcriptional level, we explored the expression profiles of a curated list of core genes involved in mitochondrial energy metabolism. This list was derived from Guarnieri et al(Guarnieri et al., 2023) to identify the potential underlying gene regulatory mechanisms with a particular focus on those that play primary roles in SL pathogenesis. Although the expression of most genes exhibited a downward trend within mitochondrial energy metabolism-associated pathways, the upregulated expression of 1–3 specific genes within crucial pathways was identified (**Figure 1C**). Specifically, pyruvate dehydrogenase kinase 1 displayed significantly elevated expression in SL skin compared to that in normal skin. This gene has previously been reported to inhibit tricarboxylic acid (TCA) cycle enzymes,(Kim et al., 2006) which may explain the relative suppression of other genes detected in the SL treatment. Moreover, we observed increased expression of enzymes implicated in glycolysis-associated metabolic pathways, such as pyruvate kinase M1/2, which is consistent with the context of aging (Abed et al., 2023). Additionally, carnitine palmitoyltransferase (CPT1A), a crucial marker of ageing-associated lipid metabolism, divulged elevated levels in SL (Chung, 2021). Insulin-like growth factor binding protein 3, a potential ageing marker that is upregulated more frequently in individuals aged over 50, was highly expressed in SL (Hong & Kim, 2018). Adenosine 5’-triphosphate binding cassette subfamily D member 1, which is associated with skin pigmentation, demonstrated a significantly increased expression in SL (Yu et al., 2022). Some genes in OXPHOS, such as cytochrome c oxidase 19 of Complex 4, also unveiled elevated expressions in SL (Kovarova et al., 2016).

### Significant alteration of fatty acid oxidation in SL interplays with energy production, redox balance, and cellular signaling

For a more comprehensive understanding of the alterations in cellular metabolism, we used a metabolic flux simulation to compute SL changes at the metabolic level. A custom-made context-specific constraint-based metabolic modelling approach, adopted and updated from our previous study, was applied (Camera et al., 2024; da Silveira et al., 2020; Guarnieri et al., 2023). Given the substantial number of flux alterations in metabolic reactions, the top 10 metabolic pathways exhibiting the most significant changes were selected and visualized in **Figure 2A**. The p-values (van der Waerden test) of the selected metabolic reactions were lower than 0.1. Among the 91 metabolic reactions demonstrating significant alterations in SL compared to the control, 39 reactions were forward and 52 were backward. As shown in **Figure 2A**, the pathways directly or indirectly involved in lipid metabolism—such as bile acid synthesis, fatty acid oxidation, glycerophospholipid metabolism, glycosphingolipid metabolism, and sphingolipid metabolism—showed significant changes, with fatty acid oxidation and mitochondrial transport displaying the most substantial alterations.

**Figure 2.**
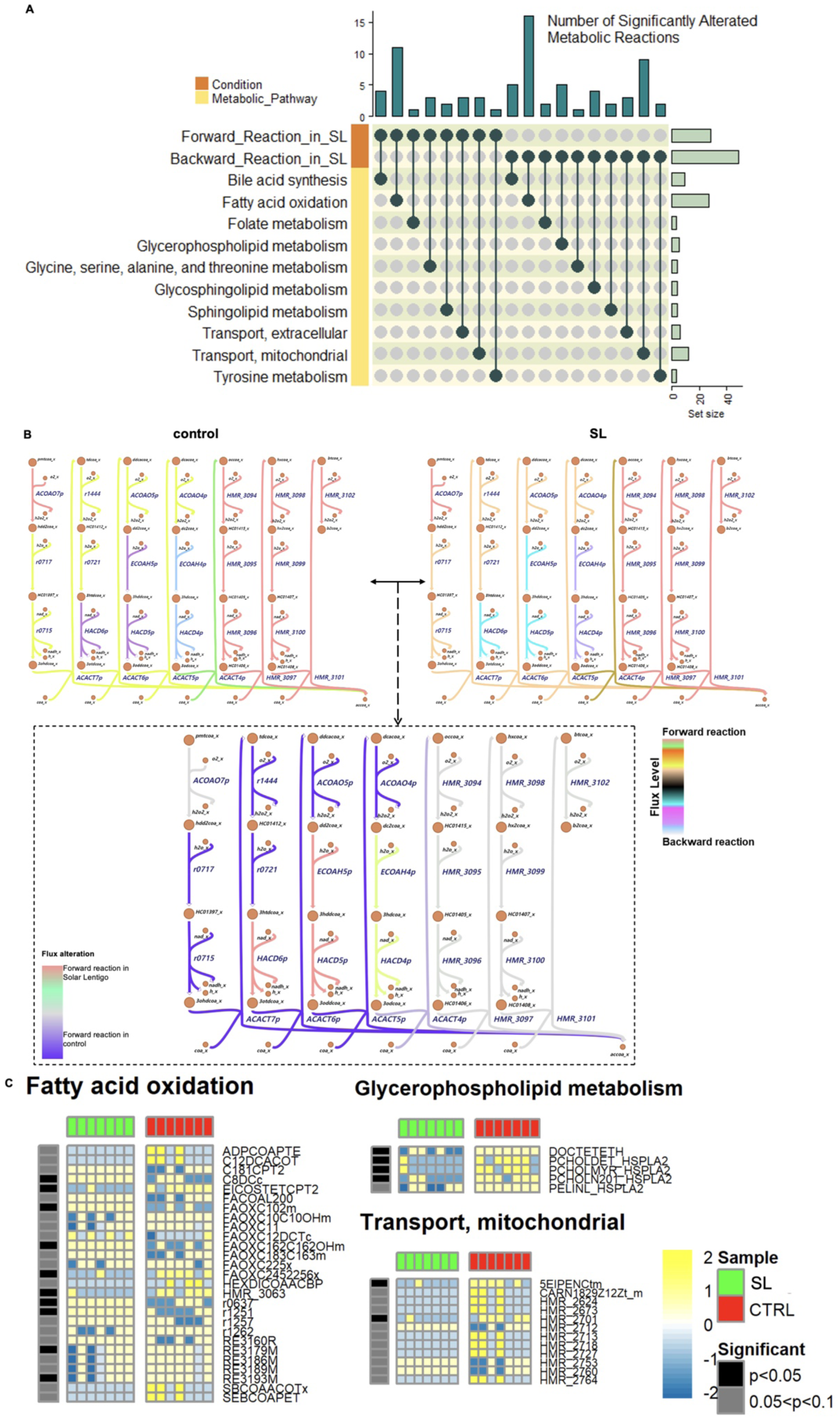
Visualisation of metabolic alterations between solar lentigo (SL) and controls using metabolic flux simulation (A) Upset plot showing the number of reactions exhibiting significant alterations *(P* < 0.1, as determined by the van der Waerden test and number of reactions > 1) between SL and controls. (B) Escher plots of fatty acid oxidation with altered reactions. Escher plots of both SL and controls presented predicted flux levels of fatty acid oxidation. The bottom plot displays the difference in levels between the two conditions. (C) Heatmaps illustrating the relative flux levels of solar lentigo and controls for three potential pathways (fatty acid oxidation, glycerophospholipid metabolism, and mitochondrial transport). Statistical significance is indicated by the leftmost bar (black: *P* < 0.05, grey: 0.05 ≤ *P* < 0.1). Heatmap colors represent row-wise Z-scores for each flux.

Given that changes in fatty acid oxidation are hypothesized to be the potential drivers of alterations in SL, we analyzed flux alterations in the core metabolic reactions of fatty acid oxidation. As depicted in **Figure 2B**, substantial forward metabolic reactions in SL were primarily NADH-yielding reactions, such as HACD6p, HACD5p, and HACD4p. Certain reactions that produce hydrogen peroxide, including ACOAO7p, HMR_3094, HMR_3098, and HMR_3102, were also slightly forward in the SL. Based on changes in the aforementioned reactions, a complex interplay between energy production, redox balance, and cellular signaling within the context of fatty acid oxidation may be involved in SL.

### Carnitine-associated reactions implicated in altered fatty acid oxidation in SL

In fatty acid oxidation, while forward reactions with considerable changes contained C8DCc (suberylcarnitine production), FAOXC102m (mitochondrial isomerization), FAOXC162C162OHm (mitochondrial fatty acid beta oxidation), and r0637 (octanoyl-CoA:L-carnitine O-octanoyltransferase), backward reactions with significant alterations included EICOSTETCPT2 (carnitine transferase), FAOXC2452256x (beta oxidation of long chain fatty acid), HMR_3063 (enoyl coenzyme A hydratase), RE3179M (3-hydroxyacyl coenzyme A dehydrogenase), and RE3193M (enoyl coenzyme A hydratase) (**Figure 2C**). Extraordinary changes in backward reactions, such as 5EIPENCtm (mitochondrial transport of 5, 8, 11, 14, 17-eicosapentenoic acid) and HMR_2701 (carnitine O-acetyltransferase), were detected in mitochondria. Taken together, considerable alterations in forward and backward metabolic reactions across fatty acid oxidation and mitochondrial transport pathways are at least partially implicated in carnitine-associated reactions, pathways, and metabolism. Carnitine is integral to cellular energy production, primarily facilitating fatty acid transport into the mitochondria (Bremer, 1983). Moreover, carnitine elevates plasma levels of adenosine triphosphate (ATP)(Capecchi et al., 1997), which is likely implicated in the transport of acetyl-CoA(Mynatt, 2009), a critical precursor in the TCA cycle (Jankowska-Kulawy et al., 2022). Therefore, alteration of carnitine-associated dynamics may lead to different energy metabolism in SL compared with that in the control.

### *CPT1B* is the potential functional gene in SL development

To identify potential functional genes in SL, we simultaneously examined molecular alterations from both transcriptional and metabolic perspectives. At the metabolic level, we focused on the computed flux differences in specific reactions between the SL and control groups. At the transcriptional level, we compared the extent of expression differences in genes predicted to encode enzymes for these reactions. Using this multi-system level analytical approach, nine metabolic reactions and eight corresponding genes were identified as potential SL markers **(Table 1)**.

**Table 1.** Biomarker candidates for solar lentigo.

| Metabolic level |  |  | Transcriptomic level |  |
| --- | --- | --- | --- | --- |
| Reaction | Primary pathway | P value (< 0.1) | Gene symbol | P value (< 0.05) |
| C8DCc<br>OCD11COACPT1_1 | Fatty acid oxidation | 0.0223<br>0.0373 | CPT1B | 0.01946217 |
| CAMPt<br>CGMPt | Transport, extracellular | 0.0373<br>0.0373 | ABCC5 | 0.043841 |
| HMR_0981 | Arachidonic acid metabolism | 0.0581 | CYP4F2<br>CYP4F3 | 0.03795117<br>0.0396098 |
| HMR_2289 | Fatty acid synthesis | 0.0581 | SCD | 0.0185095 |
| TYR3MO2 | Tyrosine metabolism | 0.0581 | TH | 0.003047084 |
| R0642 | Miscellaneous | 0.0402 | ALDH2 | 0.02901046 |
| RE3174C | Fatty acid oxidation | 0.0518 | ELOVL4 | 0.01998319 |

The candidates revealed significant alterations at both metabolic and transcriptional levels.

According to the protein-protein interaction analysis presented in **Supplementary Figure S1C**, the eight potential genes displayed a significant likelihood of interrelation under various criteria (i.e., network type, edge meaning, interaction source, and minimum interaction score). With the exception of one protein, ABCC5, two protein groups were clustered: one contained TH, ALDH2, CYP4F2, and CYP4F3 and the other included CPT1B, SCD, and ELOVL4. Some proteins not included in the potential marker list—such as CYP4A11, CYP4A22, ALOX12B, ALOX15B, and HACD4—were incorporated, but showed interrelations with each other. Five potential lists experimentally validated these interactions, indicating that they may also be functionally associated with SL.

To identify our primary target gene, which constitutes the most critical step in this study, we focused on the molecular activities that exhibited relatively large changes across multiple system levels, including the metabolic and transcriptional levels. From mathematical and computational perspectives, fatty acid oxidation-associated metabolic reactions (C8DCc and OCD11COACPT1_1) and their corresponding enzymes (CPT1B), which had the lowest p-values at the metabolic level and the third lowest p-values at the transcriptional level, were considered potential biomarkers. From a biomedical perspective, carnitine-associated molecules closely linked to energy metabolism may play a central role in altering cellular metabolism. Therefore, the fatty acid oxidation-associated enzyme CPT1B was selected for subsequent experimental validation.

### Upregulated expression of CPT1B in SL validated by immunohistochemistry

To validate the upregulation of CPT1B in the SL, immunohistochemical staining was performed on additional paired skin samples from the SL and adjacent normal skin obtained from two independent patients. CPT1B-positive cells were primarily detected in the upper dermis of both SL and control samples. CPT1B expression was visibly stronger in SL lesions than in the adjacent normal skin (**Figure 3**), which is consistent with the transcriptomic findings.

**Figure 3.**
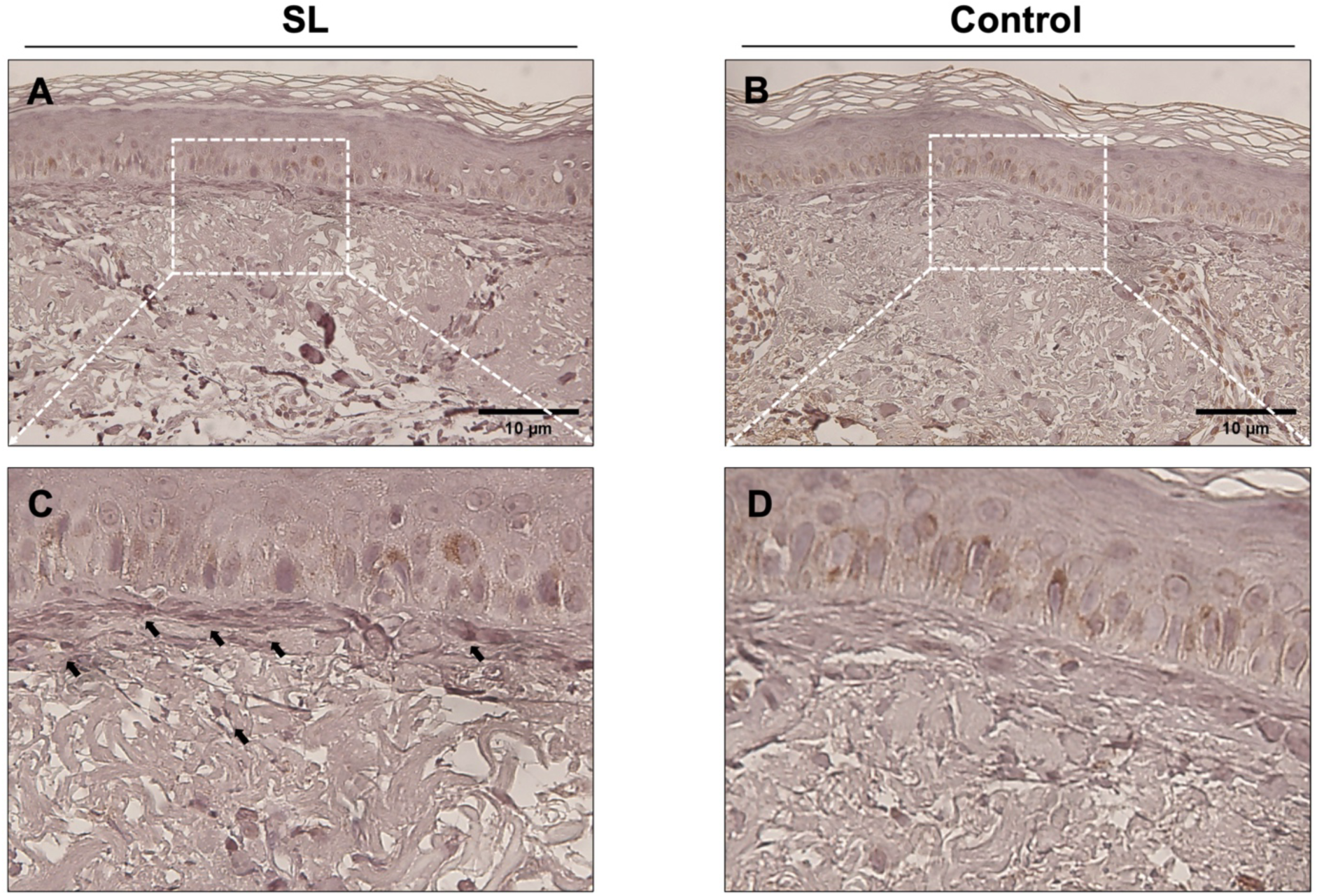
Immunohistochemical staining for CPT1B Presence of CPT1B-positive cells were found in the upper dermis. The expression of CPT1B was higher in the SL compared with control. (A, C) Solar lentigo. (B, D) Control. Black arrows indicate CPT1B-positive cells. Scale bars = 10 μm; magnification x100.

### *CPT1B* regulates pigmentation in zebrafish embryos

To explore the *in vivo* function of CPT1B in SL development, we used zebrafish embryos because the formation of these melanocytes shares many molecular patterns with those of humans (Frantz & Ceol, 2022). Zebrafish embryonic melanocytes, which originate from the neural crest, form four distinct stripes during the first three days of development: dorsal larval, lateral larval, ventral larval, and yolk sac stripes (Mort et al., 2015; Sutton et al., 2021). To examine whether CPT1B is involved in melanocyte development, we initially knocked down *CPT1B* with splicing-blocking MO targeting exon 3 and intron 3 (**Supplementary Figure S2A**). The resultant *cpt1b* morphants exhibited reduced pigmentation, particularly in the lateral larval stripe, at 3 dpf compared to the uninjected controls (**Figure 4A and 4B**). To confirm that this phenotype was specifically caused by *cpt1b* alterations, we performed a rescue experiment by co-injecting *cpt1b* MO and mRNA. Co-injection successfully restored the pigment observed in *cpt1b* morphants at 3 dpf, confirming the role of this gene in pigment formation (**Figure 4C**). However, sole overexpression of CPT1B did not affect the pigmentation (**Supplementary Figure S2B**).

**Figure 4.**
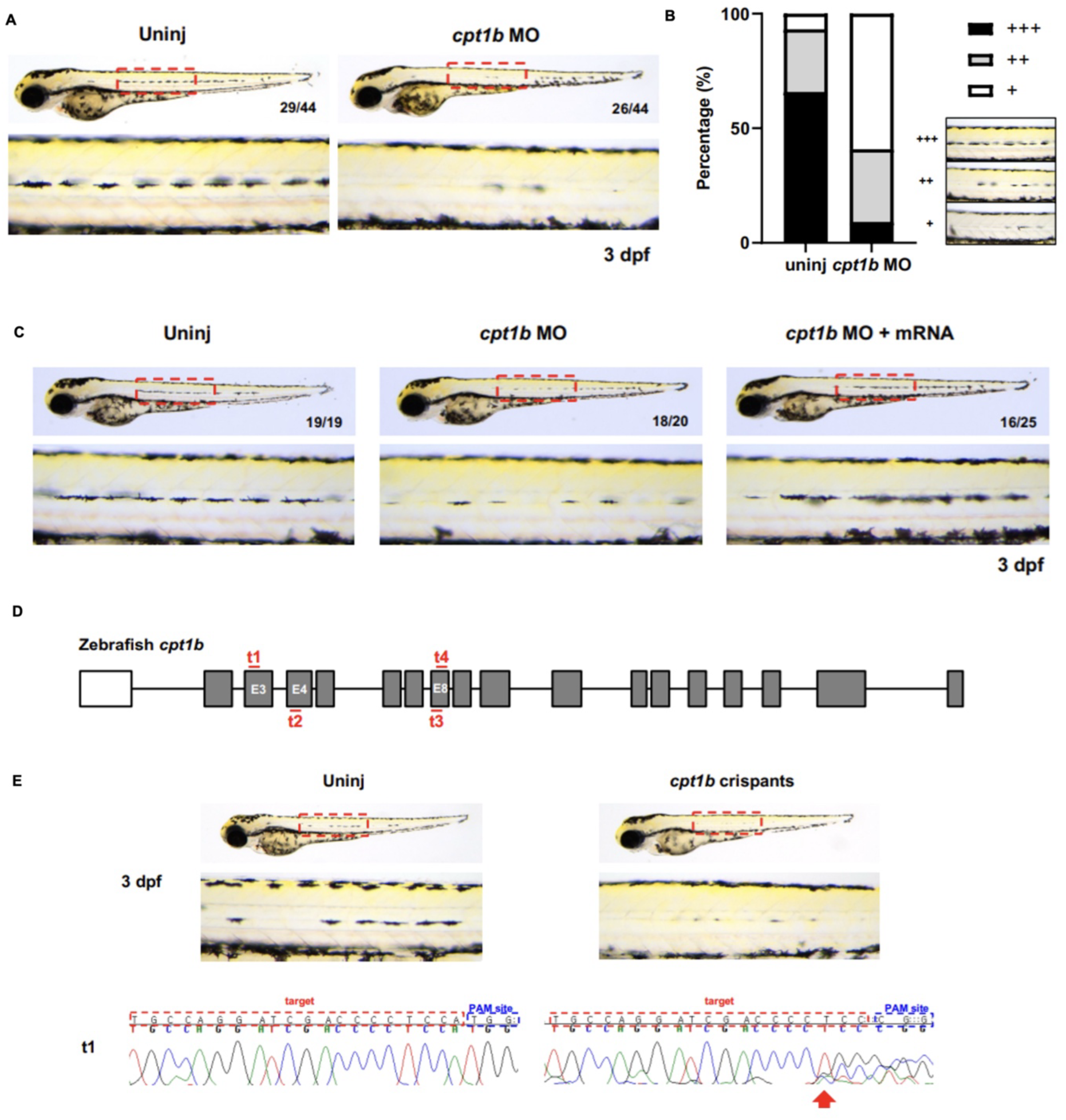
*cpt1b* is required for embryonic pigmentation (A) Representative images of live zebrafish embryos at 3 dpf: uninjected embryo (left) and *cpt1b* morphant (right). The lower panel shows a magnified view of the area within the red-dotted box on the body trunk. (B) Quantitative analysis of body pigmentation in uninjected controls and *cpt1b* morphants at 3 dpf. Pigmentation phenotypes are categorized into three groups: high (+++), middle (++), and low (+). The analysis revealed that out of 44 uninjected controls, 29 exhibited high pigmentation, 12 medium, and 3 low. In contrast, among 44 *cpt1b* morphants, 4 displayed high pigmentation, 14 medium, and 26 low. (C) Representative lateral view images of zebrafish embryos: uninjected controls (left), *cpt1b* morphants (middle), and *cpt1b* morphants with mRNA injection (right) at 3 dpf. (D) Schematic illustration of CRISPR/Cas9 targeting the zebrafish *cpt1b* gene at exons 3, 4, and 8 using four guide RNAs. (E) Representative images of zebrafish embryos at 3 dpf: uninjected controls (left) and *cpt1b* crispants (right). Below, Sanger chromatograms of DNA from an uninjected control and a *cpt1b* crispant at target site 1 are shown. The target and PAM sites are highlighted in red and blue boxes, respectively. The red arrow indicates the location of the CRISPR/cas9-mediated mutation.

The effects of CPT1B on pigmentation were explored in a knockout study using the CRISPR/Cas9 system. Previous studies have shown that CRISPR-Cas9 RNP injection resulted in high rates of biallelic mutations in G0 zebrafish, mimicking a null phenotype (Wu et al., 2018). We maximized the efficiency of obtaining biallelic mutations by injecting four guide RNAs targeting *CPT1B*, along with the Cas9 protein (**Figure 4D**). The resulting *cpt1b* crispants displayed reduced pigmentation at 3 dpf, similar to *cpt1b* morphants, compared to the uninjected controls (**Figure 4E**). Genotyping of 11 *cpt1b* crispants with reduced pigmentation revealed mutations in at least one target region in all the tested embryos (**Figure 4E and Supplementary Figure S3**). Collectively, these data indicated that CPT1B is essential for pigmentation in zebrafish embryos.

## Discussion

To adequately address a wide range of biomedical research questions with comprehensive information, recent studies have increasingly used multi-omics data acquired from multiple systems to identify novel biomarkers. Among these, the acquisition of metabolomic data has become increasingly significant, particularly for exploring alterations in cell metabolism that bridge the gap between genotype and phenotype (Fiehn, 2002). However, considering the high costs of equipment and the labor-intensive nature of experiments required to obtain metabolomics data,(Hoffmann et al., 2022) we opted for an alternative emerging analytical approach: metabolic flux modelling. Our metabolic flux simulation using Recon3D can simultaneously explore alterations in 5,835 metabolites and 10,600 metabolic reactions, which is at least three times greater than that of liquid chromatography– mass spectrometry. In the present study, we used multiple bioinformatic approaches, including metabolic modelling, to demonstrate the key molecular changes implicated in SL development at the transcriptional and metabolic levels. Significant metabolic changes related to fatty acid oxidation were found in SL, and *CPT1B* was the key regulatory gene driving the metabolic alteration in SL, which was highly upregulated compared to the control. We also validated CPT1B-induced pigmentation in knockdown (*cpt1b* MO) or knockout (*cpt1b* crispant) zebrafish embryos.

Lipids are major macromolecules with a wide range of biological functions, including energy source and storage, cellular signalling, and the membrane structure of cells and organelles. Previous studies have highlighted the roles of cellular lipid content and metabolism in cellular senescence. The enlarged morphology of senescent cells is caused by increased membrane lipids in the mitochondria and lysosomes due to the increased expression of lipogenic enzymes (Kim et al., 2010). Lipid droplet accumulation is commonly observed in senescent cells, resulting from increased lipid uptake, upregulated lipid biosynthesis, or deregulated lipid breakdown (Flor et al., 2017). In senescent cells, lipid uptake is facilitated by the increased expressions of CD36 (Saitou et al., 2018) and caveolin-1 (Linge et al., 2007) located at the cellular membrane, which could also drive cellular senescence. The synthesis of fatty acids, the building blocks of lipids, requires multiple key lipogenic enzymes, including fatty acid synthase, acetyl-CoA carboxylase, and ATP citrate lyase, which are regulated by transcription factor sterol regulatory element-binding proteins (SREBPs) (Zeng et al., 2024). Senescent fibroblasts exhibit increased expression of lipogenic enzymes as well as SREBP-1, and SREBP-1 activation could also increase cellular senescence (Kim et al., 2010).

Fatty acid oxidation is a multistep process in the mitochondria that breaks down fatty acids to generate acetyl-CoA units, which enter the TCA cycle and are linked to OXPHOS to generate energy. CPT1 is an enzyme located in the outer mitochondrial membrane that catalyzes the conversion of acyl-CoA to acylcarnitine and plays a crucial role in fatty acid oxidation (Qu et al., 2016). Ras-induced senescent fibroblasts exhibited a marked elevation in fatty acid oxidation and oxygen consumption, which was restored under the pharmacological or genetic inhibition of CPT1 (Quijano et al., 2012). CPT1 inhibition also suppressed the production of pro-inflammatory cytokines (i.e., SASPs) from Ras-induced senescent cells. The upregulation of fatty acid synthase, which is crucial for inducing cellular senescence, is also associated with increased CPT1 activity and oxygen consumption (Fafian-Labora et al., 2019). Collectively, fatty acid oxidation and CPT1 are associated with cellular senescence by modulating mitochondrial function and SASPs production.

Among the three CPT1 subtypes (CPT1A, CPT1B, and CPT1C) that exhibit tissue-specific distribution, CPT1A and CPT1B are widely distributed in humans and have considerable similarities, whereas CPT1C is exclusively present in the brain (Qu et al., 2016). The association of CPT1A expression with senescence-related diseases, such as cardiovascular disease,(Lin et al., 2022) osteoarthritis,(Jiang et al., 2022) and cancers, has been investigated;(Qu et al., 2016) however, the results were inconsistent according to cell type, indicating the tissue-specific function of CPT1A. Although several studies have reported the function of CPT1B in breast cancer,(Wang et al., 2018) type 2 diabetes mellitus,(Kim et al., 2014) obesity-related cardiomyopathy,(Zhang et al., 2016) and metabolic syndrome,(Auinger et al., 2013) its pathogenic role in skin pigmentation has not been investigated.

DNA microarray analysis of SL samples showed higher expression of fatty acid metabolism-related genes, especially fatty acid desaturases and arachidonate 15-lipoxygenase, than normal sun-exposed skin (Aoki et al., 2007). Fatty acids regulate melanocyte activity and melanogenesis via proteasomal degradation of tyrosinase (Ando et al., 2004). In a temporal analysis of B16 melanocytes where the transition from basal depigmented to pigmented state occurs slowly over 6 days, metabolic changes occurred with increased fatty acid oxidation during the melanogenic phase that exhibits the peak melanogenic activity. αMSH-induced activation of SREBP1 drives the metabolic programming, and inhibition of SREBP1 causes decreased pigmentation and tyrosinase activity (Sultan et al., 2022). Collectively, although evidence is lacking on the association between CPT1B and pigmentation, altered fatty acid metabolism, which is highly correlated with cellular senescence, can affect melanogenesis and pigmentation in SL as shown in the present study.

Zebrafish are valuable models for melanocyte and human melanoma research because many fundamental molecular mechanisms are shared between humans and zebrafish, facilitating the study of pigment regulation across various pigmentary phenotypes (Mort et al., 2015; Pickart et al., 2004). For example, screening using the *mitfa* zebrafish mutant in a melanoma-prone Tg(*mitfa*:BRAFV600E); *p53-/-* background has confirmed that GDF6, a BMP ligand, significantly impacts melanoma formation (Ceol et al., 2011; Venkatesan et al., 2018). Moreover, studies have shown that patients with albinism possessing the *C10orf11* nonsense mutation exhibit reduced melanocyte numbers and pigmentation, a finding supported by zebrafish models (Gronskov et al., 2013). Consequently, zebrafish are suitable model organisms for studying various human pigmentation disorders.

## Conclusion

Compared with paired adjacent normal skin, SL revealed basal hyperpigmentation, solar elastosis, dermal melanophages, and rete ridge elongation. Exploring the interactions between cellular senescence and metabolic alterations provides a new perspective for explaining the development of age-related diseases. Herein, we demonstrate the transcriptional-metabolic alteration processes implicated in SL development by identifying the corresponding regulatory mechanisms of differentially activated metabolic reactions and their potential enzymatic genes at the multisystem level using multiple bioinformatics approaches. Upregulated expression of CPT1B was confirmed in an immunohistochemical study and was validated to regulate pigmentation in a zebrafish model. Targeting CPT1B and lipid metabolism may offer novel therapeutic strategies for the management of SL and age-related pigmentation.

## Data availability statement

The data presented in this study are available upon request from the corresponding author.

## Ethics statement

This study was conducted in accordance with the Declaration of Helsinki and the International Conference on Harmonization and Good Clinical Practice Guidelines and was reviewed and approved by the Institutional Review Board of Kyung Hee University Hospital at Gangdong (KHNMC 2022-04-014).

## Author Contributions

Conceptualization, Yoonsung Lee, Man S Kim, and Soon-Hyo Kwon;

Investigation, Yoonsung Lee, Man S Kim, and Soon-Hyo Kwon;

Formal Analysis, Yueun Choi and Jihan Kim;

Methodology, Uijeong Nam and Hae-Na Moon;

Software, Soo-Young Yoon., Hanseul Cho, and Seoyeon Kim;

Visualization, Sanghyeon Yu and Jihan Kim;

Resources, Bark-Lynn Lew and Soon-Hyo Kwon;

Supervision, Yoonsung Lee, Man S Kim, and Soon-Hyo Kwon;

Writing—original draft preparation, All authors;

Writing—review and editing, Yoonsung Lee, Man S Kim, and Soon-Hyo Kwon;

All authors have read and agreed to the published version of the manuscript.

