## Supplementary for "CPT1B-Mediated Fatty Acid Oxidation Induces Pigmentation in Solar Lentigo"

### Supplementary materials

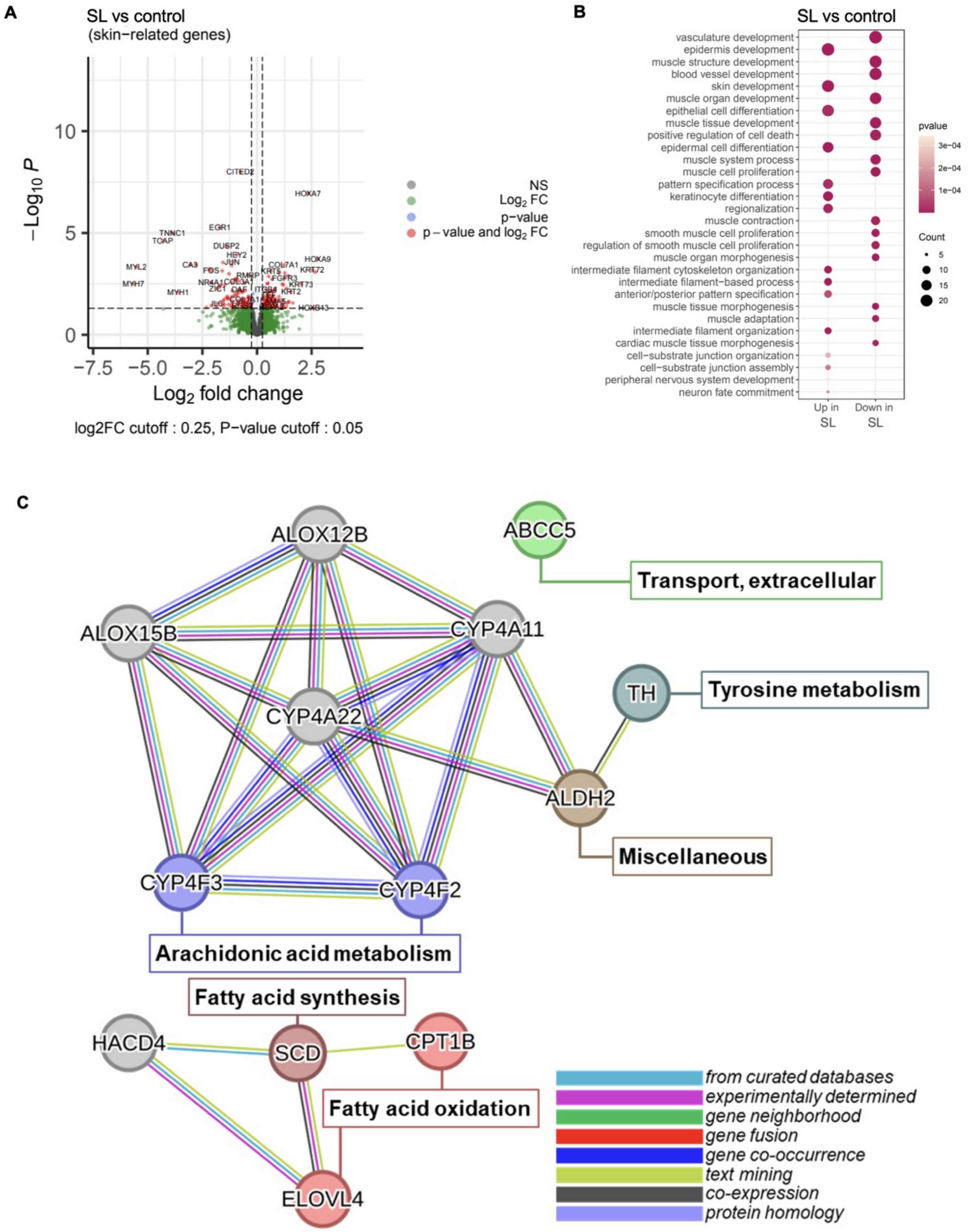


**Supplementary Figure S1.** Differential expression, enrichment pathway analysis and protein-protein interaction network

(A) Volcano plot with differentially expressed genes between SL and controls, highlighting those reaching statistical significance using red dots *(P* < 0.05 and |log2FoldChange| > 0.25). (B) Dot plot displaying top 15 enriched gene ontology pathways identified based on the differentially expressed genes from A *(P* < 0.05). (C) Protein-protein interaction network constructed using STRING database, showing interconnectedness between selected proteins. The connections were based on evidence from multiple databases, highlighting their validity and significance.

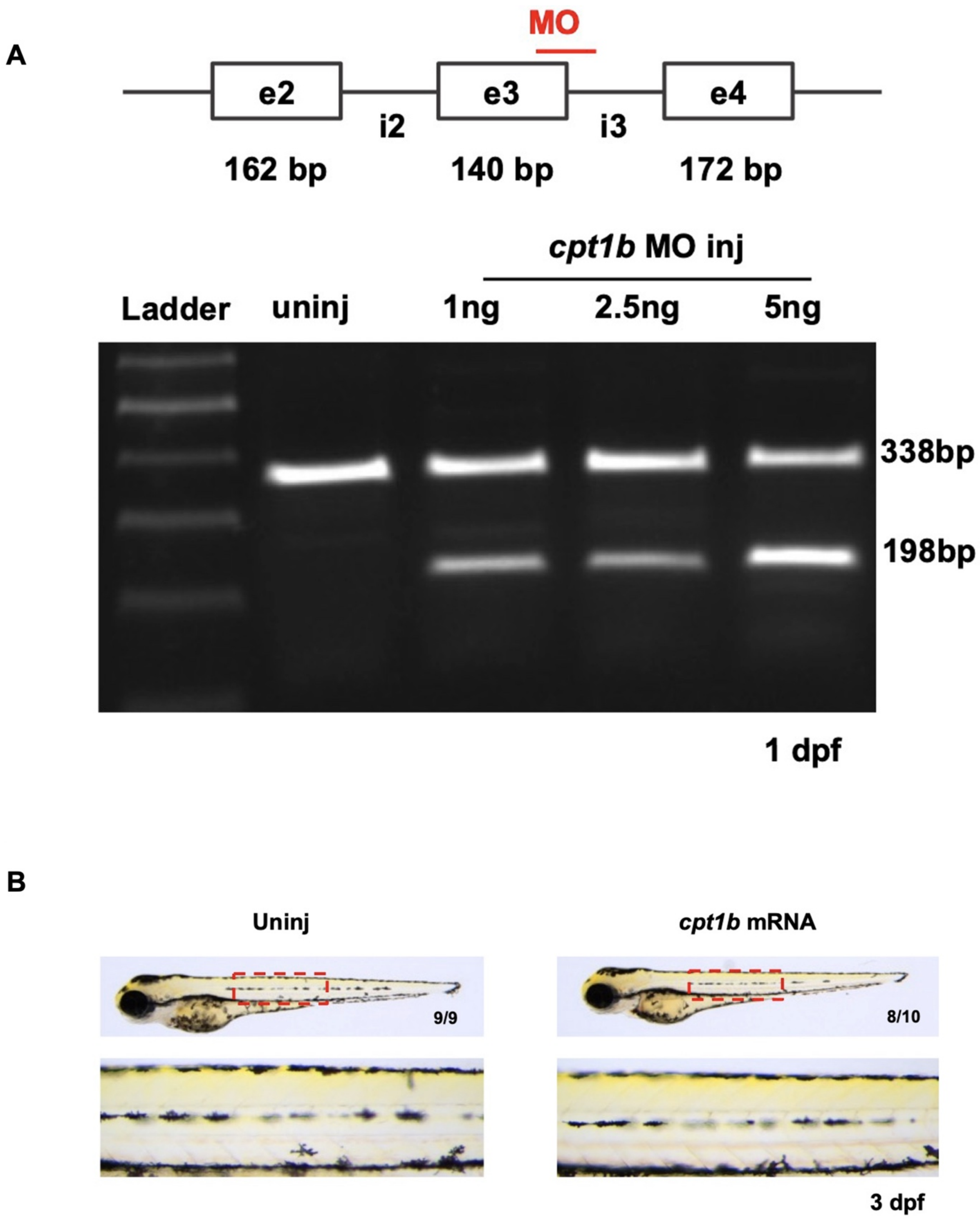


**Supplementary Figure S2.** *cpt1b* knockdown using a splice-blocking morpholino (MO) and *cpt1b* overexpression via mRNA injection.

**(A)** Schematic diagram showing the targeting of exon 3 (e3) and intron 3 (i3) of the zebrafish *cpt1b* gene by a splice-blocking MO. The accompanying gel electrophoresis image displays, from left to right: a DNA ladder (lane 1), uninjected controls (lane 2), and *cpt1b* morphants (lanes 3–5) injected with 1, 2.5, and 5 ng of cpt1b MO, respectively. Compared to the control, exon 3 skipping was observed in the morphants, indicating successful splicing inhibition.

**(B)** Representative live images of uninjected (left) and *cpt1b* mRNA-injected (right) embryos at 3 dpf. The lower panels show a magnified view of the trunk region (highlighted by a red box), illustrating pigmentation differences.

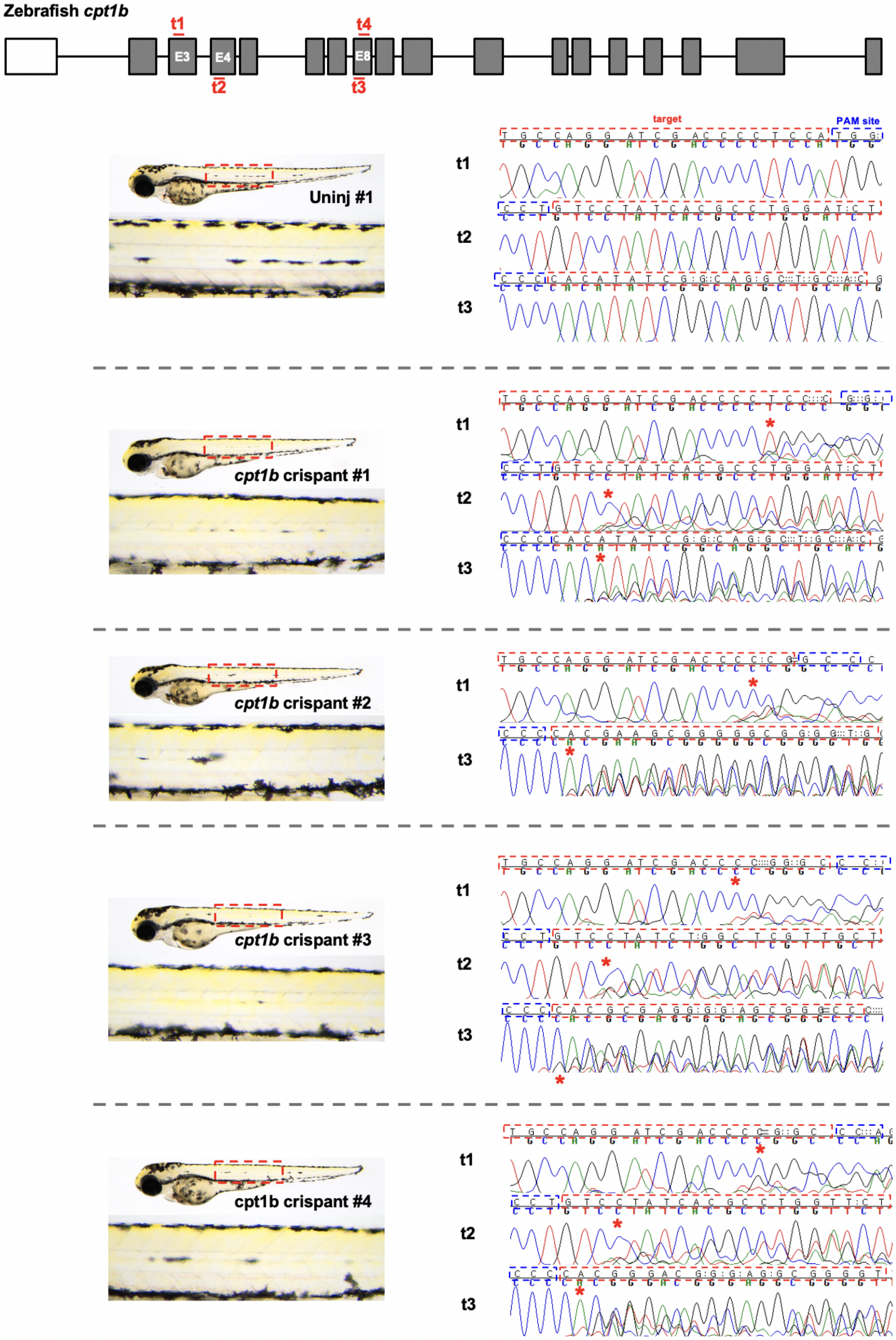
Zebrafish cpt1b
t1
t4
E3
E4
E8
t2
t3
target
PAM site
Uninj #1
t1
t2
t3
cpt1b crispant #1
t1
t2
t3
cpt1b crispant #2
t1
t3
cpt1b crispant #3
t1
t2
t3
cpt1b crispant #4
t1
t2
t3

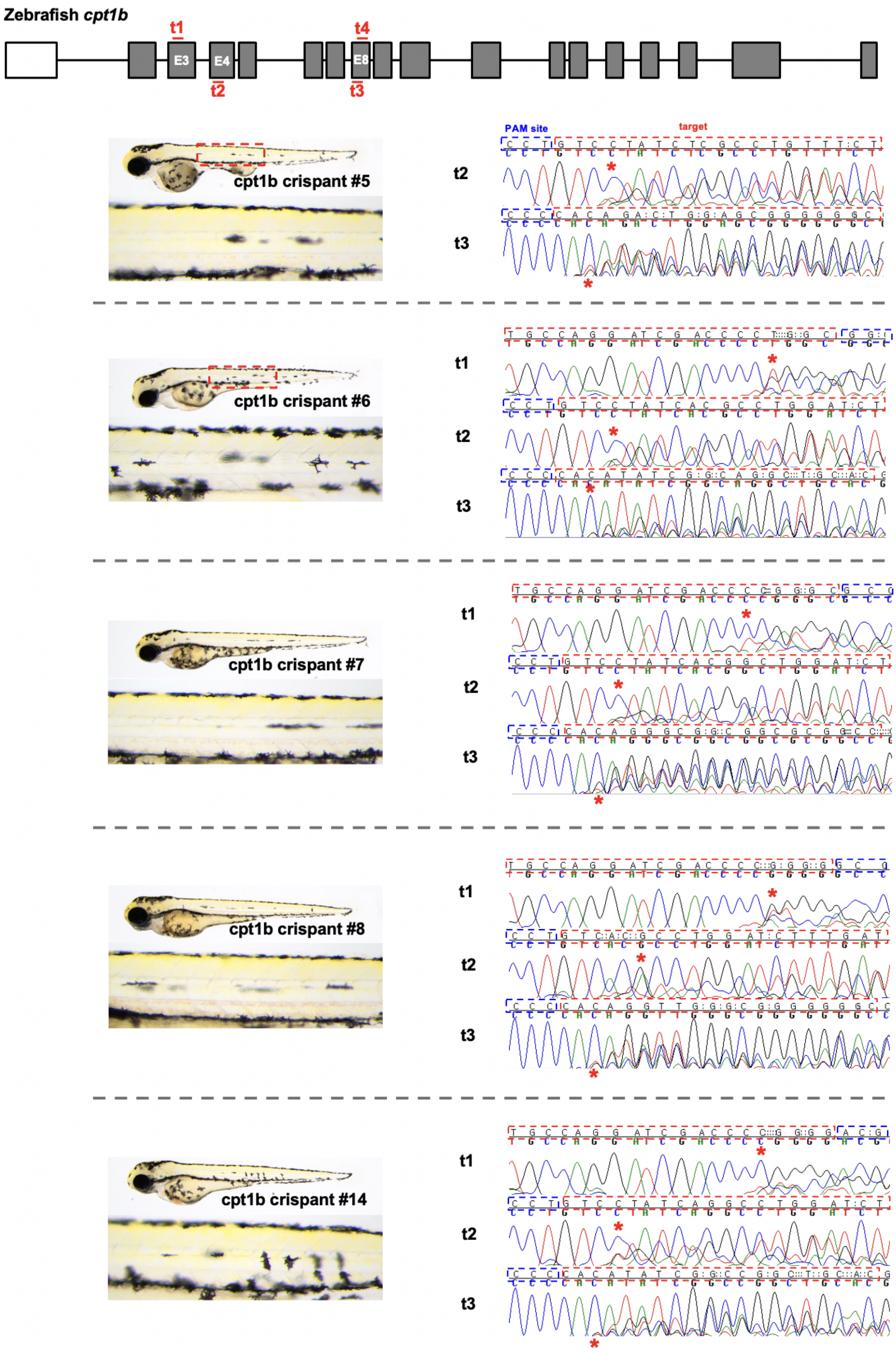
Zebrafish cpt1b
t1
t2
t3
t4
E3
E4
E8
PAM site
target
cpt1b crispant #5
cpt1b crispant #6
cpt1b crispant #7
cpt1b crispant #8
cpt1b crispant #14

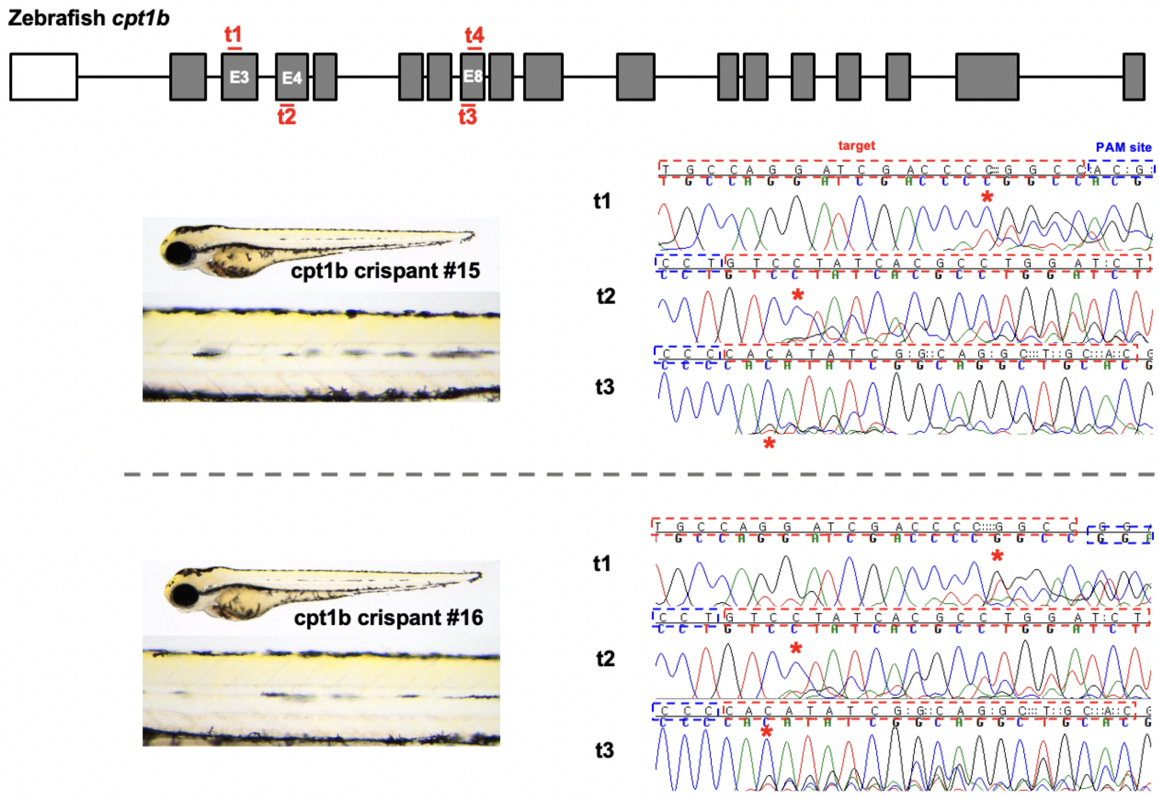


**Supplementary Figure S3.** Sanger sequencing chromatograms of DNA at target sites of uninjected control and *cpt1b* crispant zebrafish embryos

The schematic illustrates the locations targeted by the *cpt1b* guide RNAs, corresponding to exons 3, 4, and 8. The left panel shows images of one uninjected control and 11 *cpt1b* crispants exhibiting reduced pigmentation phenotypes among 16 total crispants. The right panel displays Sanger chromatograms for the embryo shown on the left. Only chromatograms with confirmed mutations are presented. Red and blue dotted boxes indicate the target and PAM sites, respectively. Red asterisks mark the positions of CRISPR/Cas9-induced mutations.
